# Bayesian Neural Networks for Cellular Image Classification and Uncertainty Analysis

**DOI:** 10.1101/824862

**Authors:** Giacomo Deodato, Christopher Ball, Xian Zhang

## Abstract

Over the last decades, deep learning models have rapidly gained popularity for their ability to achieve state-of-the-art performances in different inference settings. Novel domains of application define a new set of requirements that transcend accurate predictions and depend on uncertainty measures. The aims of this study are to implement Bayesian neural networks and use the corresponding uncertainty estimates to improve predictions and perform dataset analysis. We identify two main advantages in modeling the predictive uncertainty of deep neural networks performing classification tasks. The first is the possibility to discard highly uncertain predictions to increase model accuracy. The second is the identification of unfamiliar patterns in the data that correspond to outliers in the model representation of the training data distribution. Such outliers can be further characterized as either corrupted observations or data belonging to different domains. Both advantages are well demonstrated on benchmark datasets. Furthermore we apply the Bayesian approach to a biomedical imaging dataset where cancer cells are treated with diverse drugs, and show how one can increase classification accuracy and identify noise in the ground truth labels with uncertainty analysis.

## 1. Introduction

Deep neural networks have seen a dramatic increase in popularity in recent years, due to their outstanding performances in complex prediction tasks (Krizhevsky et al., 2012; LeCun et al., 2015). The main drawback of neural networks lies in their lack of interpretability (they are often deemed as “black boxes” (Benítez et al., 1997; Shrikumar et al., 2017; Lundberg and Lee, 2017)) and their dependence on point estimates of their parameters. Despite their ability to outperform simpler models, a single prediction score (i.e. the accuracy of the prediction) is not sufficient for a variety of tasks and domain applications (Ghahramani, 2015). Modeling applications such as healthcare require an additional feature to the prediction score, that is a measure of confidence that reflects the uncertainty of the predictions. For example, a neural network performing diagnosis of brain tumors by analyzing magnetic resonance images needs a way to express the ambiguity of an image in the same way as a doctor may express uncertainty and ask for experts help. Moreover, predictive uncertainty provides further insights about the data because more certain predictions correspond to cleaner data both from a technical and a contextual point of view.

The output of neural networks is often treated as a probability distribution. However, despite there is a correlation between the accuracy of the prediction and this confidence score, that is the output value, this should not lead to think that it is an appropriate measure of uncertainty as this would show that the model makes mostly overconfident predictions (Gal and Ghahramani, 2016).

Instead of the described prediction score, we analyzed data by means of the predictive uncertainty, that can be decomposed into epistemic uncertainty, which stems from model’s parameters as well as the specific architecture of the model, and aleatoric uncertainty, which depends on the noise of the observations (Der Kiureghian and Ditlevsen, 2009).

In the remaining sections of this paper we present our implementation of Bayesian neural networks using variational inference and our confidence measure formulation (2), we briefly analyze the most relevant alternative to our approach (3) and we show our results over multiple datasets belonging to different domains (4). Finally, we discuss the results and the advantages of our approach (5).

## 2. Methods

The idea behind Bayesian modeling is to make predictions by considering all possible values for the parameters of the model. In order to do so, the parameters are treated as random variables whose distribution is such that the most likely values are also the most probable ones. The posterior distribution of the model parameters, conditioned on the training data, is defined using Bayes theorem:

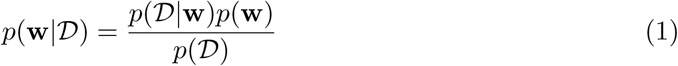

where **w** is the set of model parameters and ***𝒟*** is the training set. The estimation of the posterior requires the definition of a likelihood function *p* (***𝒟***|**w**), a prior density *p* (**w**), and the computation of the marginal likelihood *p* (***𝒟***) that is unfeasible for complex models such as neural networks.

### 2.1. Mean Field Variational Inference

Variational inference allows to avoid computing the marginal likelihood by directly approximating the posterior distribution with a simpler one (Jordan et al., 1999). In order to do so, it is necessary to minimize the Kullback–Leibler (KL) divergence between the proposed distribution and the posterior. The KL divergence is defined as follows:

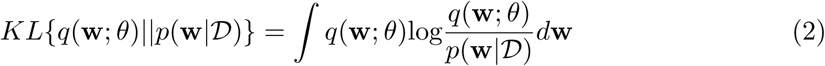

where *θ* is the set of variational parameters describing the proposed distribution *q* of the model’s parameters **w**. Since the posterior distribution is not known, we need to define a different objective to minimize the KL divergence. Such objective function is called Evidence Lower Bound (ELBO) and it is defined as follows:

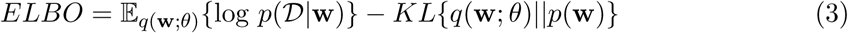

Variational inference turns the marginal likelihood computation problem into an optimization one: maximizing the ELBO as a function of the variational parameters so that the proposed distribution fits the posterior (proof in Appendix A).

We approximated the posterior with a multivariate Gaussian distribution and, in order to simplify the optimization process, we used the mean field approximation. This choice allowed us to avoid a quadratic increase of the parameters to optimize by factorizing the posterior approximation:

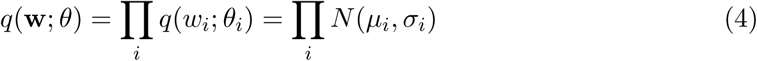

### 2.2. Bayesian Neural Network Training

In order to be able to apply the backpropagation algorithm to the variational parameters of a Bayesian neural network, we applied the local reparameterization trick (Kingma et al., 2015) which separates the deterministic and the stochastic components of the weights, which are random variables. Furthermore, we also reparameterized the standard deviations of the weights using the softplus function to keep them positive.

The loss function of the Bayesian neural network is the variational objective, i.e. the negative ELBO, where the likelihood can be divided in the sum of the contributions of all the individual data points in the dataset (Hoffman et al., 2013; Mandt et al., 2016, 2017) and it is possible to employ a minibatch approach (Graves, 2011; Blundell et al., 2015):

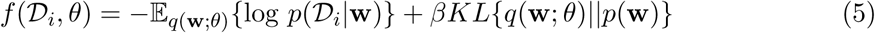

where ***𝒟***_*i*_ represents the *i*-th mini-batch, *β* = 1 */M* is the scaling factor of the KL divergence due to the minibatch approach, and *M* is the number of mini-batches. The prior distribution is a Gaussian distribution that is equivalent to L2 regularization (Blundell et al., 2015).

### 2.3. Predictive Uncertainty

The probability distribution over the parameters of the model yields a predictive distribution whose mean can be approximated using Monte Carlo samples from the posterior approximation:

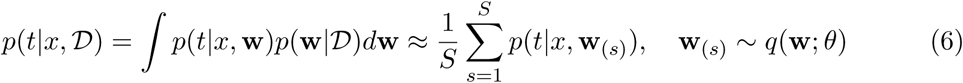

where *S* is the number of weights samples taken, therefore the number of output samples too. The corresponding predictive uncertainty can be computed starting from the variance of the predictive distribution, considering both the epistemic and aleatoric components of uncertainty (Kwon et al., 2018; Kendall and Gal, 2017):

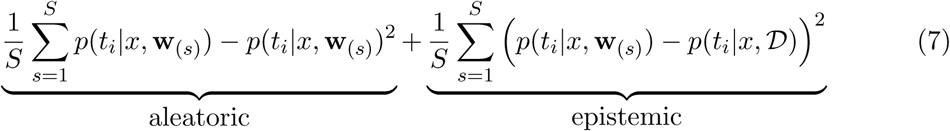

where *i* is the index of the predicted class. Finally, we used the function [INEQ] to transform such uncertainty measure into a more intuitive Bayesian confidence score which is also easier to compare to the standard neural networks confidence score.

## 3. Related work

There exist many solutions to find the posterior distribution of complex models such as Bayesian neural networks. Among them, it is worth citing the work regarding the Laplace approximation (Ritter et al., 2018), that approximates the posterior with a Gaussian distribution, and the Markov Chain Monte Carlo methods (Neal et al., 2011), that approximate the posterior by directly sampling from it. The most relevant of such techniques is dropout (Hinton et al., 2012), a regularization technique that, when active both at training and test time, has been proved to be equivalent to the approximation of a Bayesian neural network (Gal and Ghahramani, 2016).

Dropout has been extensively used to approximate Bayesian neural networks because of its ease of implementation and retrieval of predictive uncertainty estimates. Moreover, dropout has shorter training and prediction times when compared to the previously mentioned approaches. For these reasons, dropout based Bayesian uncertainty measures have also been used to perform biomedical image analysis and prediction (Dürr et al., 2018; Leibig et al., 2017). However recent work has exposed the main limitations of such approach, mainly related to the use of improper priors and the correctness of the variational objective (Hron et al., 2018).

## 4. Results

In this section, we compare the Bayesian approach and the standard one on the MNIST dataset (LeCun and Cortes, 1998), we analyze the predictive uncertainty to find out-of-distribution data from a closely related dataset (EMNIST by Cohen et al.(2017)), and we validate our method against cellular microscopy images. The architectures of the neural networks and the corresponding training hyperparameters are discussed in Appendix C.

### 4.1. Standard and Bayesian neural networks comparison

We first trained a Bayesian convolutional neural network (LeNet5) using the MNIST training set and tested the model on the corresponding test set with *S* = 100 output samples (see Appendix B for discussion of the number of predictive samples). As demonstrated by predictions for 200 example images (Figure 1(*a*)), some images have a consistent prediction, while others produce more than one classification result over the range of samples, which suggests low confidence. Overall, predictions performed using the output samples individually showed good classification accuracy with small fluctuations (Figure 1(*b*)). For each image, we computed the Bayesian prediction by taking the average of the output samples as explained in the Methods section and illustrated in Figure 1(*c*)and Figure 1(*d*).

**Figure 1:**
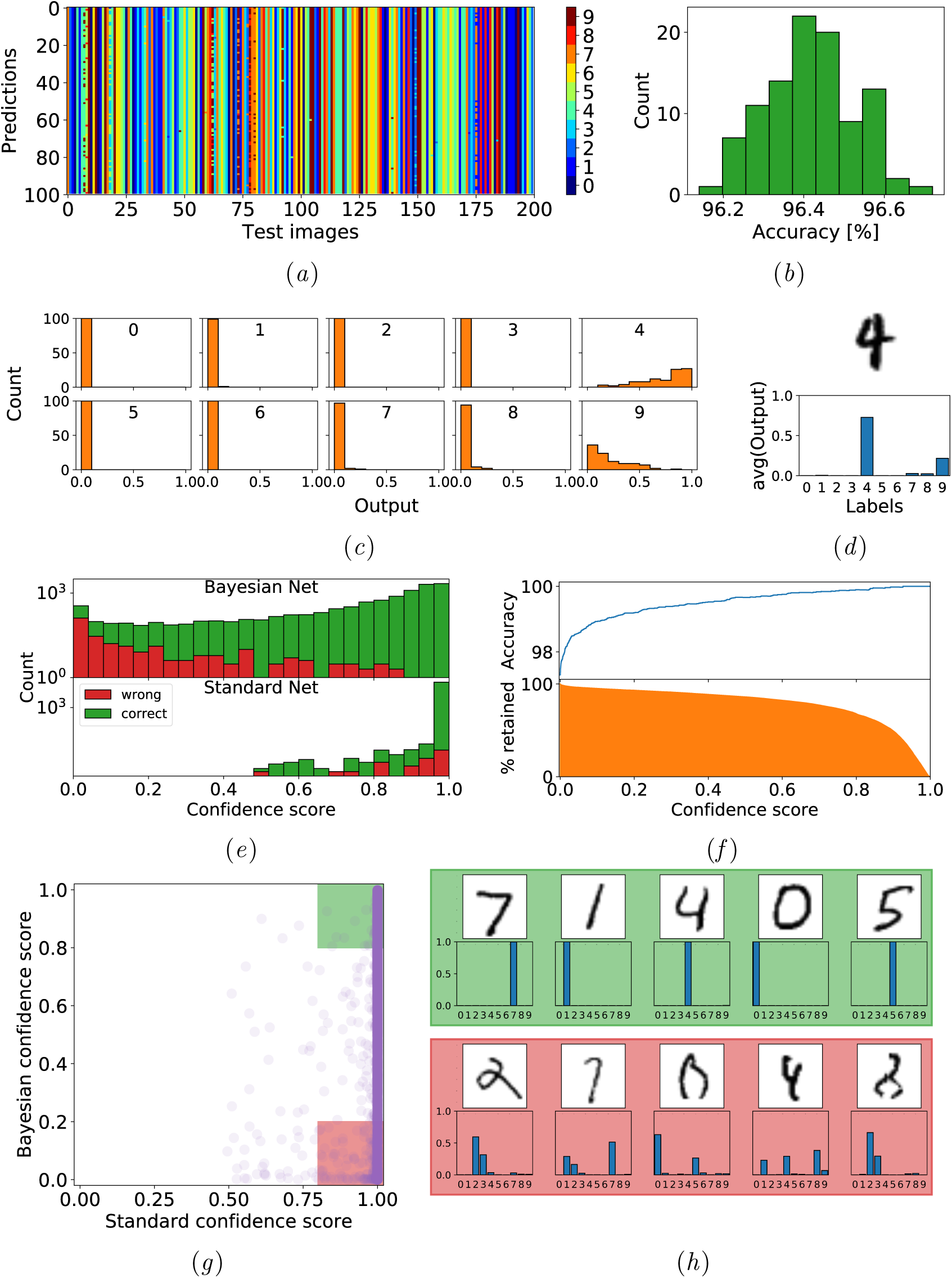
(a) Predictions from 100 output samples of 200 test images; (b) Corresponding histogram of accuracy; (c) Per class histograms of 100 output samples; (d) Corresponding final output for a test image of an ambiguous 4; (e) Confidence score distribution over the MNIST test set; (f) Increasing Bayesian confidence score cutoff increases accuracy while decreases the percentage of retained images; (g) Comparison of Bayesian and standard confidence scores over the MNIST test set; (h) Image samples, with related outputs, predicted with the corresponding colored areas confidence scores.

We then compared the Bayesian prediction results with a standard neural network trained on the same MNIST data. As shown in Figure 1(*e*), confidence scores from the standard neural network tend to be close to 1 and there is no differentiation between correct and wrong predictions. In contrast, Bayesian confidence scores are high for correct predictions and low for incorrect ones. Therefore it is possible to improve the overall accuracy by increasing the confidence threshold and retaining only high-confidence predictions as illustrated in Figure 1(*f*)). We also plotted the confidence scores from the Bayesian neural network and the standard neural network against each other for each test image (Figure 1(*g*)). By examining individual images in Figure 1(*h*), we see that images with high Bayesian confidence are the canonical digits while images with low Bayesian confidence correspond to corrupted or ambiguous observations. The standard neural network however is not able to distinguish them.

### 4.2. Out of distribution data

The EMNIST dataset is designed with the same image format as MNIST, but it expands to include hand-written letters. In order to validate the capability of the model to identify out-of-distribution samples — that is, images that cannot be labeled with any of the possible classes — we performed predictions over the EMNIST dataset with the Bayesian neural network previously trained on the MNIST dataset. As shown in Figure 2(*a*), the model predicts most numbers with high confidence, while it predicts letters with low confidence as they do not belong to the domain of the MNIST training data. We examined the confidence score distribution of each letter and illustrated in Figure 2(*b*)three representative examples. As expected, the letter “o” is predicted as 0 with high confidence while the letter “w”, that does not resemble any of the digits, is predicted with low confidence. Interestingly, the confidence scores of letter “i”, show a bimodal distribution. After manually checking individual images, we realized that some “i”s are very similar to the number 1 and are predicted with high confidence, while other “i”s are written with a dot on top, therefore considered as out-of-distribution samples with low confidence predictions.

**Figure 2:**
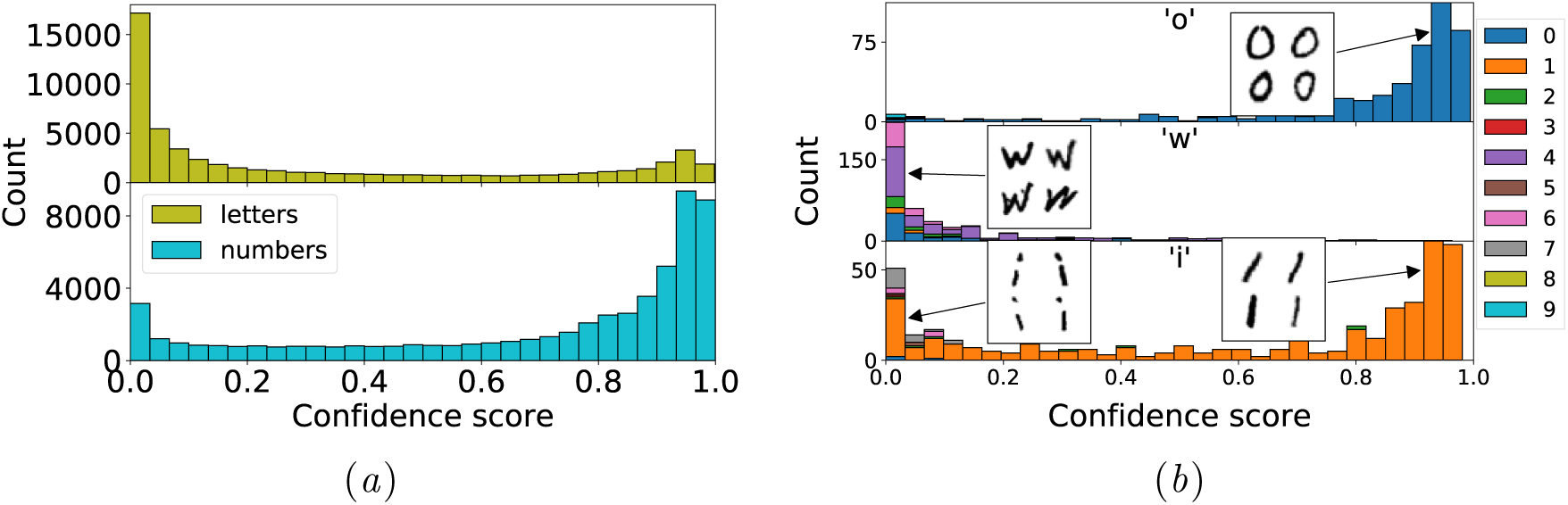
(a) Confidence score distribution of letters and numbers in EMNIST; (b) Three example letters: “o”, “w” and “i”.

### 4.3. Cellular microscopy images

As part of the Broad Bioimage Benchmark Collection (BBBC), the BBBC021 dataset is made of microscopy images of human MCF-7 breast cancer cells treated with 113 compounds over eight concentrations and labeled with fluorescent markers for DNA, F-actin, and *β*-tubulin (Ljosa et al.,2012; Caie et al., 2010). A subset of BBBC021 with 38 compounds is annotated with a known mechanism of action (MoA) and it has been widely used to benchmark diverse analysis methods (Ljosa et al., 2013; Kandaswamy et al., 2016; Godinez et al.,2017; Ando et al., 2017). The MoA annotations come both from visual inspection by experts as well as scientific literature, using a course-grained set of 12 MoA, with each MoA containing multiple compounds and concentrations.

We applied the Bayesian approach, as described above, to a simplified version of the Multi-Scale Convolutional Neural Network (MSCNN) previous designed by Godinez et al. (2017). For validation, we adopted the rigorous leave-one-compound-out process, where in each session, all except one compound are used to train the model and the hold-out compound is used for validation. Figure 3(*a*)illustrates the predictive confidence for all hold-out compounds. As expected, the wrong predictions distribution has a peak on the low confidence side while the correct predictions distribution has a peak on the high confidence side. Thus with increasing threshold, one can improve overall classification accuracy as shown in Figure 6(*a*)Appendix D.

**Figure 3:**
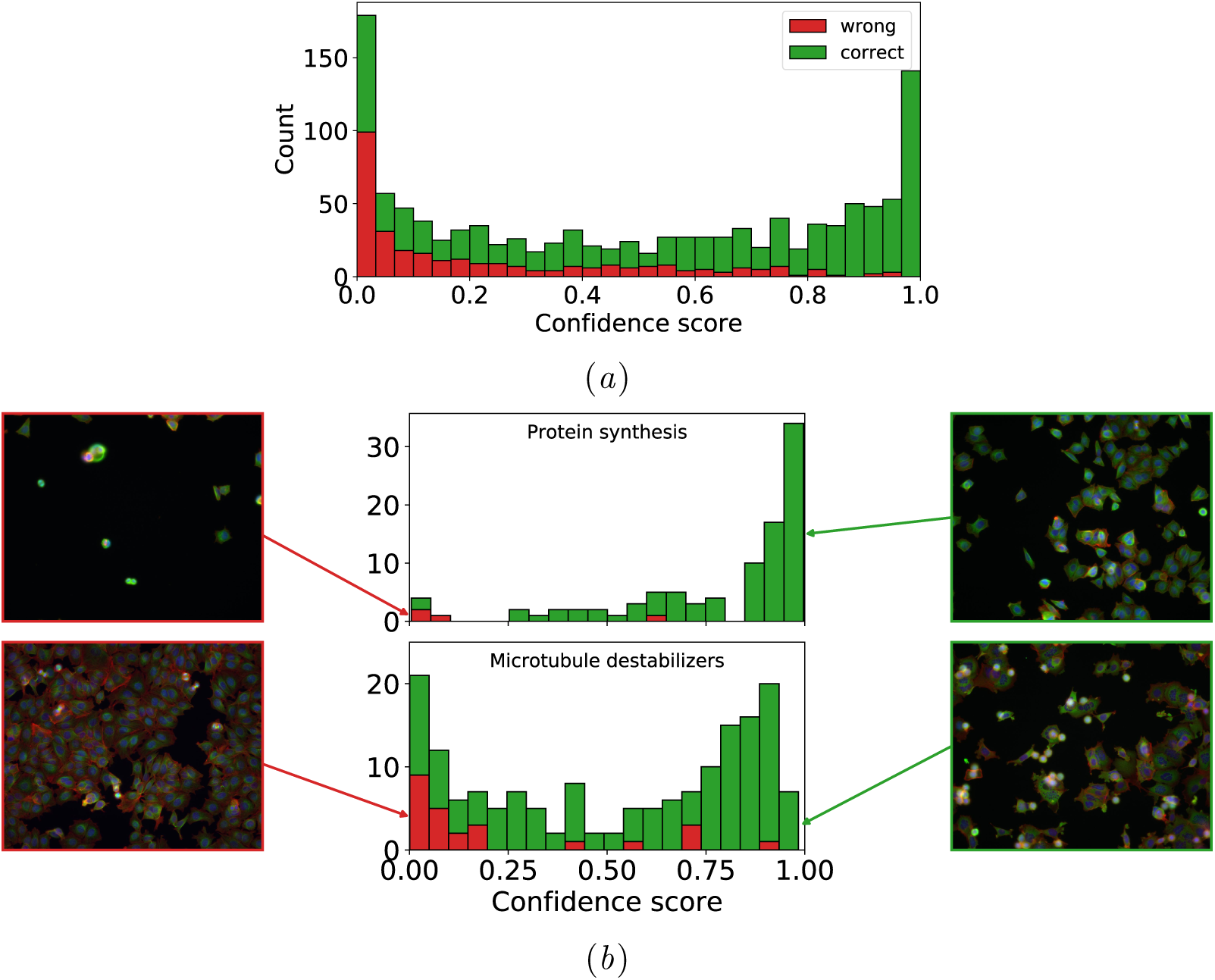
(a) Confidence score distribution of all hold-out compounds from BBBC021; (b) Two representative classes: protein synthesis and microtubule destabilizers, and the corresponding example images.

The effects of compound treatment are complex due to how compounds interact with one or multiple protein targets and how these interactions are reflected on cell morphology labeled with fluorescent markers. For this reason, we further examined the Bayesian confidence scores for each of the 12 MoA and displayed two of the most relevant ones in Figure 3(*b*).

Observations belonging to the “protein synthesis” MoA (96 images and three compounds) are mainly predicted correctly. As expected, the confidence score of the four images predicted incorrectly is 0.04 *±* 0.05 (median *±* median absolute deviation), substantially lower than the correctly predicted images at 0.92 *±* 0.10. Moreover, such images are considerably different that the rest, in fact they are mostly black, probably due to noise during the acquisition and annotation processes.

Similarly, incorrect predictions of “microtubule destabilizers”, show a different morphology than the expected one. These anomalous images correspond to cells treated with colchicine, one of the 4 compounds associated with this MoA (see Figure 6(*b*)in Appendix D). This observation is consistent with the results of unsupervised approaches on the same dataset (Godinez et al., 2018) and indicates an error in the ground truth annotations. Furthermore, most of the 168 images of microtubule destabilizers are predicted correctly but some of them have low confidence scores. This is due to the corrupted observations from colchicine, that are used to train the model in order to validate the other compounds.

## 5. Discussion

The performance of deep neural networks in computer vision tasks has been explored for biological research and drug development. Compared with natural images, biomedical images have various challenges, such as noisy labels and out-of-distribution samples. To address these challenges, we have implemented a Bayesian neural network with variational inference and exploited the confidence score derived from the predictive variance of the model. Such uncertainty measure proved to be more precise than the standard neural networks confidence score on simple, well known, benchmark datasets, as well as complex biomedical ones. Moreover, we were able to identify multiple anomalies in cellular images and the corresponding annotations, therefore we believe this Bayesian neural network approach has added value to the field of biomedical image classification.

Ground-truth labels for biomedical images are often impossible or prohibitively expensive to generate, which is why the field has moved towards unsupervised approaches. We envision future applications of the Bayesian framework to clustering and unlabeled images retrieval.

## Acknowledgments

GD would like to acknowledge Prof. Maurizio Filippone (Eurecom, Department of Data Science) for the supervision of GD’s Master thesis^1^ from which this work derives.

## Footnotes

1. https://webthesis.biblio.polito.it/10920/1/tesi.pdf

## Appendix A. Evidence Lower Bound derivation

Variational inference employs the exclusive version of the KL divergence. Since the posterior distribution is not known, a different objective needs to be defined starting from the logarithm of the marginal likelihood of the model:

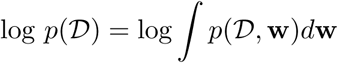

The computation of the marginal likelihood is the core issue of Bayes theorem, in order to proceed we use an auxiliary function which corresponds to the proposal for the posterior approximation *q* (**w** *|θ*) that we write as *q* (**w**) to simplify the notation:

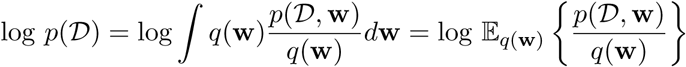

Then, it is needed to bring the logarithm inside the expectation which can be done by applying the Jensen’s inequality:

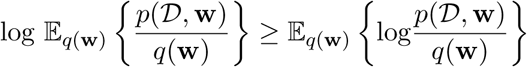

Since we are now using an inequality, its right term is a lower bound of the logarithm of the marginal likelihood, also called model evidence, hence the name Evidence Lower Bound (ELBO). The ELBO can now be reformulated in the following way:

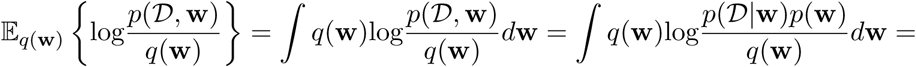

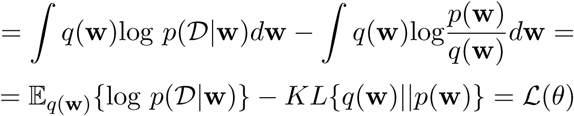

Maximizing this lower bound with respect to the variational parameters *θ* of *q* (**w**; *θ*) provides a value as close as possible to the logarithm of the marginal likelihood and it is equivalent to minimizing the initial KL divergence between *q* (**w**) and *p* (**w**|***𝒟***):

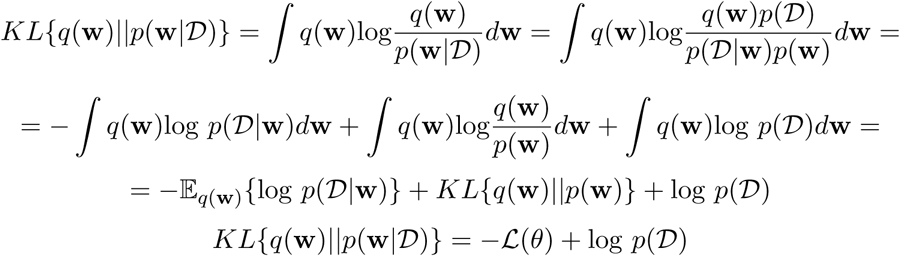

The marginal likelihood is constant, therefore maximizing the ELBO is equivalent to minimizing the KL divergence between the posterior and its approximation.

## Appendix B. Number of predictive samples

The predictive distribution is approximated using Monte Carlo samples, therefore, depending on the number of outputs averaged to compute the final prediction, the corresponding accuracy can be more or less precise. Figure 4 shows 100 accuracies per numbers of samples used to compute the output, and the corresponding median and variance per number of samples. As the number of samples increases we get a closer estimate of the real accuracy value and its variance decreases. Figure 4 data correspond to a Bayesian neural network trained on the MNIST dataset.

**Figure 4:**
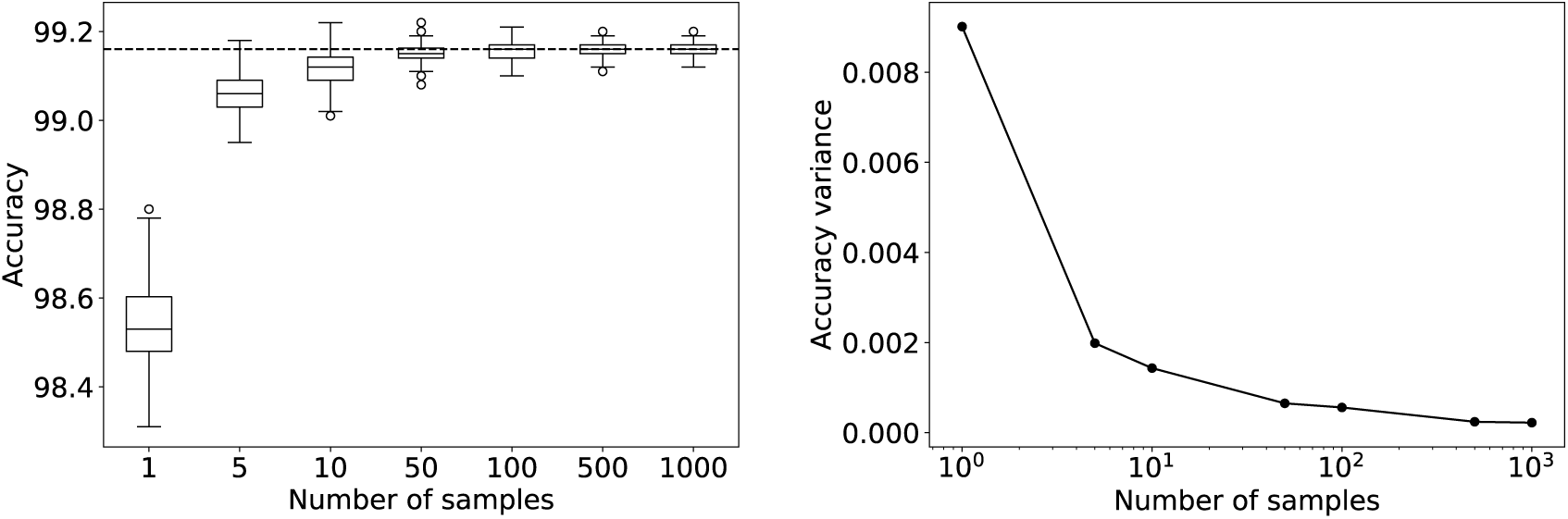
(left) Boxplot of 100 values of the accuracy of a Bayesian neural network per different number of samples and (right) the variance per number of samples. The dashed line corresponds to the median of the accuracies with 1000 samples.

## Appendix C. Neural networks architectures and training hyperparameters

The architectures used for the experiments are illustrated in Figure 5. We used the same architecture (LeNet 5) for both the standard and Bayesian case in order to be consistent when performing comparison. We performed the optimization of the neural networks parameters using the Adam optimizer (Kingma and Ba, 2014). The training hyperparameters for all the architectures used are illustrated in Table 1.

**Table 1:** Training hyperparameters of the different architectures used in the experiments.

| Neural network architecture | Dataset | Learning rate | Batch size | # Epochs |
| --- | --- | --- | --- | --- |
| Standard LeNet5 | MNIST | $10^{-3}$ | 256 | 200 |
| Bayesian LeNet5 | MNIST | $10^{-3}$ | 32 | 300 |
| Bayesian MSCNN | BBBC021 | $10^{-2}$ | 20 | 80 |

**Figure 5:**
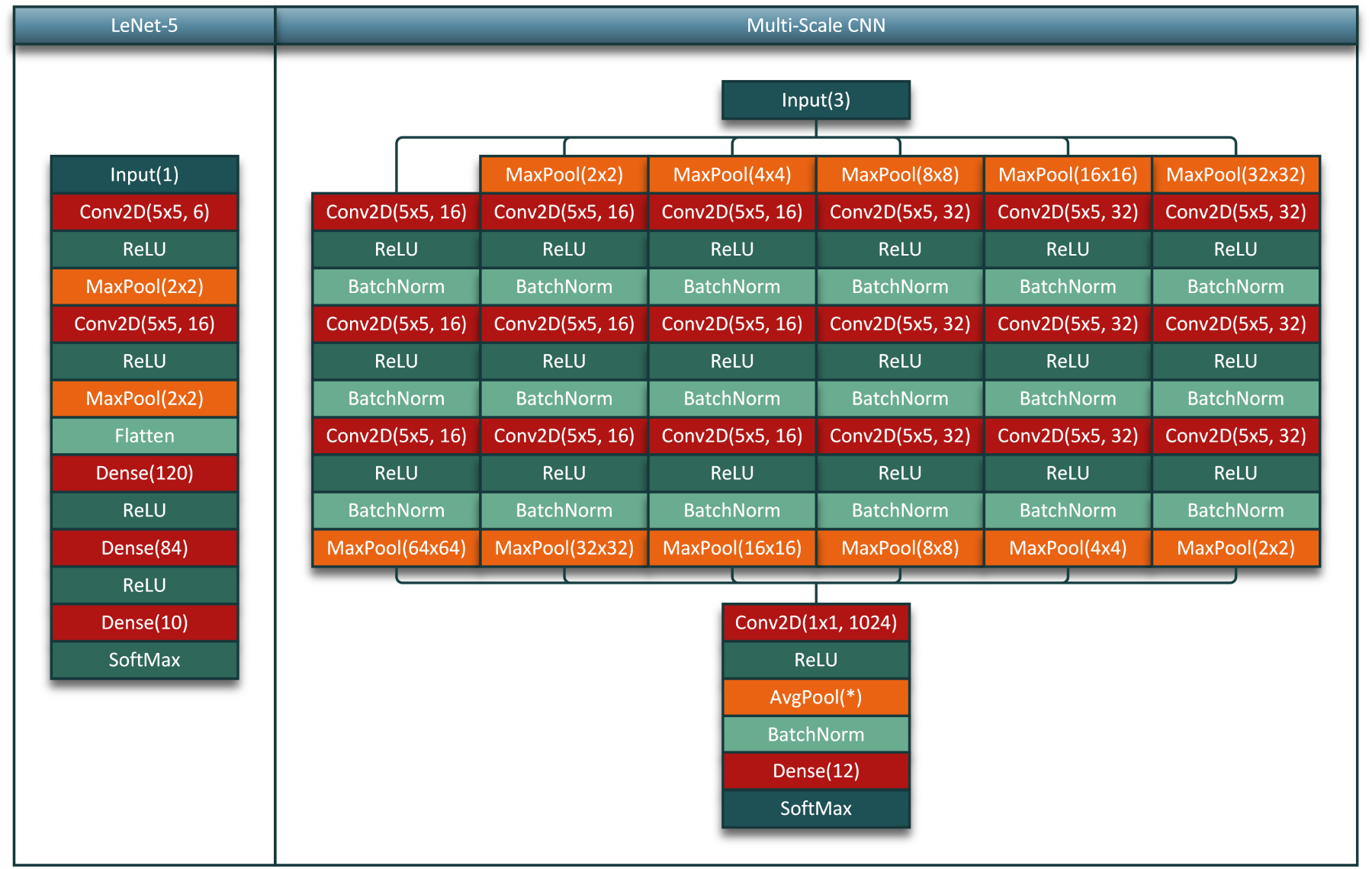
Neural networks architectures.

In order to train the standard LeNet5 we used the cross entropy loss function and a learning rate schedule consisting of a single step at epoch 170, decreasing the learning rate by a factor of 10. Furthermore, we added weight decay with a factor of 5 *×* 10^*−*3^. It is important to underline that this is not the best model to train on the MNIST dataset, this implementation choice has been taken because the model is still able to perform very well and, since the dataset is contextually simple, further improvements could lead to higher accuracies but complicate the comparison with the Bayesian counterpart.

Given the different nature of Bayesian LeNet5 with respect to the standard version, it has been trained with a lower batch size and for more epochs, with a single learning rate step at 200 epochs. The loss function is the variational objective, namely the ELBO loss. No weight decay has been applied because the regularization is already imposed by the shape and parameters of the prior.

In order to train the MSCNN, we adopted a weighted loss function because of the BBBC021 classes imbalance. The weights of this loss were computed as the inverse of the number of observations for each class, so that all the classes have the same influence on the updates of the model parameters. We finally trained the neural network using a learning rate schedule consisting of a single step at epoch 60.

## Appendix D. BBBC021 analysis

**Figure 6:**
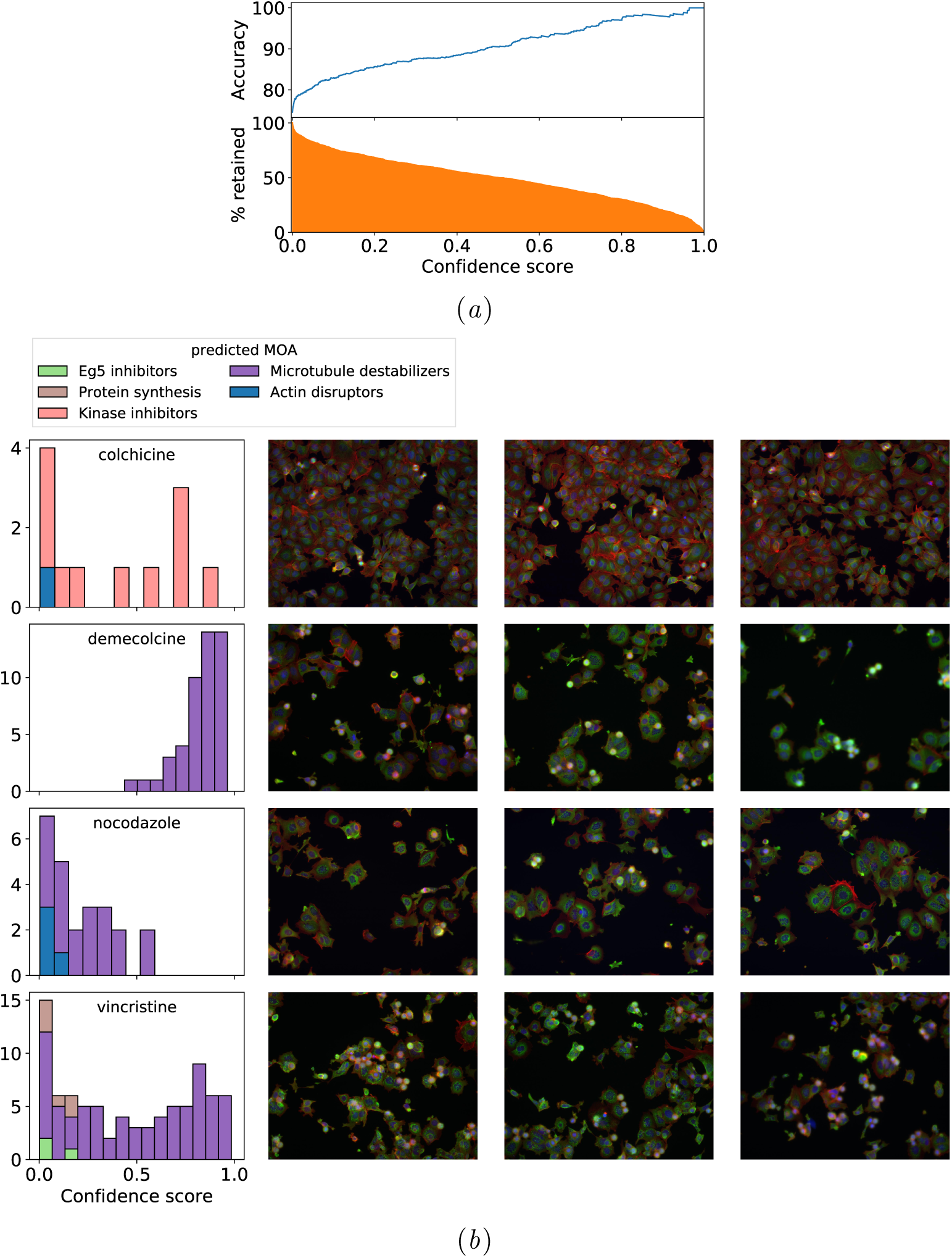
(a) Confidence score threshold analysis for BBBC021 predictions (same as Figure 1(*f*); (b) Confidence score distribution of each compound belonging to the microtubule destabilizers MoA and corresponding image samples.

